# Translational readthrough goes unseen by natural selection

**DOI:** 10.1101/844621

**Authors:** April Snofrid Kleppe, Erich Bornberg-Bauer

## Abstract

Occasionally during protein synthesis, the ribosome bypasses the stop codon and continues translation to the next stop codon in frame. This error is called translational readthrough (TR). Earlier research suggest that TR is a relatively common error, in several taxa, yet the evolutionary relevance of this translational error is still unclear. By analysing ribosome profiling data, we have conducted species comparisons between yeasts to infer conservation of TR between orthologs. Moreover, we infer the evolutionary rate of error prone and canonically translated proteins to deduct differential selective pressure. We find that about 40% of error prone proteins in *Schizosaccharomyces pombe* do not have any orthologs in *Saccharomyces cerevisiae*, but that 60% of error prone proteins in *S. pombe* are undergoing canonical translation in *S. cerevisiae*. Error prone proteins tend to have a higher GC-content in the 3’-UTR, unlike their canonically translated ortholog. We do not find the same trends for GC-content of the CDS. We discuss the role of 3’-UTR and GC-content regarding translational readthrough. Moreover, we find that there is neither selective pressure against or for TR. We suggest that TR is a near-neutral error that goes unseen by natural selection. We speculate that TR yield neutral protein isoforms that are not being purged. We suggest that isoforms, yielded by TR, increase proteomic diversity in the cell, which is readily available upon sudden environmental shifts and which therefore may become adaptive.

**Author Summary:** There is an evolutionary balance act between adaptation and selection against change. Any system needs to be able to adapt facing novel environmental conditions. Simultaneously, biological systems are under selection to maintain fitness and thus undergo selection against mutations. Phenotypic mutations - translational errors during protein synthesis - have been suggested to play a role in protein evolvability by enabling quick assessment of viable phenotypes and thus enable quick adaptation. Here we test this hypothesis, by inferring evolutionary rate of proteins prone to a specific case of phenotypic mutations: translational readthrough (TR). By making use of publicly available data of yeasts, we find that TR goes unseen by natural selection and appear as a neutral event. We suggest that TR goes unseen by selection and occurs as “permissive wallflowers”, which may become relevant and yield adaptive benefits. This work highlights that stochastic processes are not necessarily under stringent selection but may prevail. In conclusion, we suggest that TR is a neutral non-adaptive process that can yield adaptive benefits.

## Introduction

During protein synthesis, ribosoxmes translate mRNA into proteins. Translation synthesis is terminated when the ribosome encounters a stop codon in an mRNA. Occasionally, stop codons are ignored by the ribosome, and translation continues beyond the coding sequence (CDS), into the untranslated region (3’-UTR). This process, called translational readthrough (TR), yields a protein chain that becomes longer than one would predict from the DNA sequence alone. Alterations to the molecular phenotype of a protein that are not encoded in the DNA, but are introduced during transcription or translation, are referred to as phenotypic mutations. Phenotypic mutations are also known as translational and transcriptional errors. Translational and transcriptional error rate is manifold higher than the genotypic error rate [1–3], i.e. the error rate during DNA replication, which leads to heritable variation. Phenotypic mutations have been suggested to act as intermediate stepping stones for traits not yet encoded on the DNA level [4]. In the context of two correlated point mutations, which are jointly required for the formation of stabilising interactions in a protein, this has lead to the formulation of “the look-ahead effect” [4]. Assuming only one of the two point mutations are present on DNA level, and the second mutation is expressed by a phenotypic mutation, the look-ahead effect hypothesis explores the probability of the fixation of a novel phenotype prior to genotypic encoding. If the novel phenotype has a significantly higher fitness than the already encoded protein, both the phenotypic mutation and the first point mutation will get fixed in the population. Once the phenotypic mutation is fixed, Whitehead et al. Whitehead calculated that the probability for encoding the phenotypic alteration on DNA level is highly probable. Accordingly, phenotypic mutations are suggested to enable exploration of protein sequences, which enables quick assessment of viable phenotypes. Quick assessment of viable phenotypes will in turn yield a higher evolvability. Analogously, erroneously translated proteins are predicted to have a high evolutionary rate. Prior analysis of the look-ahead effect was strictly theoretical and computational, while controlling for population size. However, in recent years experiments indicate that phenotypic mutations may become adaptive. A study in bacteria by Bratulic et al. [5] found that phenotypic mutations are foremost selected to be tolerated and not purged, which supports the assumption that phenotypic mutations may be tolerated and at some point become adaptable. Studies in fungi displayed experimental evidence for how phenotypic mutations may yield not only a fitness effect [6–8], but appear prior to gene duplication and then become encoded [6]. In light of these studies, we hypothesize that proteins prone to TR should have a higher evolutionary rate if they are yielding beneficial phenotypes. Previously, we have investigated TR in yeast, to identify protein features that correlate with TR-rate. We identified high gene expression to be the most prevalent feature, next to highly disordered C-termini [9]. However, whether these features are a consistent trait in the presence of TR, remains untested. Here, we wanted to test our previous speculations regarding what features may buffer or coincide with TR, and whether one can trace TR - as a phenotypic mutation - to high evolvability. By analysing publicly available ribosome profiling data, we identify proteins that undergo TR and cluster these proteins as “ *leaky*”. Proteins undergoing canonical translation are clustered as “ *non-leaky*”. This is done both for *Schizosaccharomyces pombe* and *Saccharomyces cerevisiae*, and we have here analysed structural features regarding *leaky* proteins and their orthologs. Furthermore, we treat TR as a phenotypic mutation and investigate the look-ahead effect’s prediction; whether proteins undergoing TR have an elevated evolutionary rate, and whether TR show signs of being adaptive. We have investigated this by analysing evolutionary rate for proteins in *S. cerevisiae*, and compared the ratio of proteins undergoing positive selection between *leaky* and *non-leaky* proteins.

## Results and Discussion

### Inferring translational readthrough

We analysed ribosome profiling data for *S. cerevisiae* [10] and for *S. pombe* [11]. By analysing expression data of wildtype cells from an enriched medium from [10] and [11], we inferred translational readthrough (TR) that occur unrelated to external stress factors. We identified proteins that undergo TR and clustered these proteins into what we refer to as a *leaky* set. Proteins that undergo canonical translation, we clustered as the *non-leaky* set. The clustering of *leaky* and *non-leaky* sets were conducted individually for each investigated species, as in [9] and is described in detail in Materials and Methods. We found 80 *leaky* proteins and 3200 *non-leaky* proteins for *S. cerevisiae*. For *S. pombe*, there were 240 identified *leaky* proteins, and 2830 *non-leaky* proteins. We compared our sets to the data from our previous inference of TR in *S. cerevisiae* [9]. We found an overlap of 30 proteins between the *leaky* sets. There were no *leaky* proteins that overlapped with *non-leaky* sets, in either data set across the two inferences. We infer that the lack of complete overlaps, between the *leaky* data, is due to variance of gene expression across the ribosome profiling studies.

### Ortholog comparisons

We wanted to investigate TR across species. The aim was primarily to explore whether *leaky* proteins have orthologs, and whether such ortholog proteins are *leaky* too. A consistent pattern of TR across orthologs could indicate that TR is an intrinsic feature to the protein coding sequence. Alternatively, if *leaky* proteins have *non-leaky* orthologs, comparative analyses of sequence features may reveal what enables or prevents TR. Moreover, identifying sequence features may elucidate the evolutionary path between error-prone versus canonical translation. In order to analyse homologous of *leaky* proteins, we aimed to infer orthologs between *S. cerevisiae* and *S. pombe*, as there is accessible ribosome profiling data, in addition to that these species have well annotated genomes. We compared *leaky* and *non-leaky* orthologs between *S. cerevisiae* and *S. pombe*. We find that only 3% of *leaky* proteins in *S. pombe* have a *leaky* ortholog in *S. cerevisiae*. Totally 58% of *leaky* proteins, and 64% of *non-leaky* proteins in *S. pombe*, have at least one *non-leaky* ortholog in *S. cerevisiae*. This suggest that TR is not conserved, at least not between these two species. With some single exceptions (see Table S1), the remaining *leaky* proteins in *S. pombe* do not have any orthologs in *S. cerevisiae*.

In a previous study in yeasts by Yanagida et al. [6], it was found that TR yielded functional protein isoforms, which maintained organismal fitness under stressed conditions. These isoforms were near identical to paralogs in closely related species that had undergone whole genome duplication (WGD). Yanagida et al [6] suggested that TR yield functional isoforms, prior to being encoded in the DNA, in support of the look-ahead effect hypothesis. Given the study of Yanagida et al. [6], we wondered if we would find functionally encoded 3’-UTR in *S. pombe*, who did not undergo a WGD. In order to infer whether *leaky* protein of *S. pombe* yield an isoform similar to their *non-leaky* ortholog in *S. cerevisiae*, we conducted two kind of alignments. First, the protein sequence of the *non-leaky* orthologs in *S. cerevisiae* were aligned to their respective *leaky* ortholog CDS sequence, of *S. pombe*. We used an exhaustive bestfit-model in exonerate [12] to align the sequences (see Materials and Methods). Secondly, we conducted the same alignment, but we included the 3’-UTR to the CDS of *leaky* sequence. This allowed us to infer whether the alignment score of orthologs increased when including the 3’-UTR. If the 3’-UTR is under neutral selection and foremost influenced by mutation bias, the alignment score should be lower when including the 3’-UTR. Alternatively, the alignment should result in a higher alignment score when including the 3’-UTR, if the 3’-UTR has a high sequence similarity to the orthologous protein. Indeed, we found this to be the case for 12 genes of 43 aligned genes (27%). In other words, a portion of isoforms yielded by TR in *S. pombe* is encoded in the CDS of *S. cerevisiae*. This result may imply that the common ancestor lost a stop codon in *S. cerevisiae* and the sequences have undergone convergent evolution [13–15]. Alternatively, a stop codon may have been introduced in *S. pombe* and the 3’-UTR has been under purifying selection in *S. pombe*. Regardless of the evolutionary path responsible for the homology between the given 3’-UTRs and CDS termini, the homology is only present in a subset of genes. The majority of ortholog alignments does not reveal a 3’-UTR homology to the CDS, which suggest that the 3’-UTRs are generally not conserved past species divergence. We therefore find it unlikely that the given subset of isoforms would be homologous by chance. We speculate whether the isoforms in question may be both functional and under selection. Moreover, our results suggest that the findings of Yanagida et al. [6] is probably not an isolated single-case for the particular protein they investigated. In conclusion, we have demonstrated that TR is not conserved across orthologs in these two species, and also that proteins undergoing TR are not purged. Moreover, a subset of *leaky* isoforms yielded by TR are highly homologous to their encoded *non-leaky* orthologs. This finding suggests that TR may yield functional isoforms, and that the 3’-UTR may be under selection for coding amino acids.

### Protein feature analysis

Next to identifying homologous genes, we wanted to deduct what features may impact or coincide with TR. First, we compared features for *leaky* and *non-leaky* sets within same-species, Fig S1 and Table S2 and S3. Data for *S. pombe* and *S. cerevisiae* can be found in S3File.csv and S4File.csv, respectively. Secondly, we compared orthologous genes that differ with respect to TR, by comparing *leaky* and *non-leaky* proteins in *S. pombe* with their *non-leaky* ortholog in *S. cerevisiae*. When investigating the orthologs, we used the subtracted difference for each feature. For example, if we wanted to investigate gene expression for a protein in *S. pombe*, we subtracted the gene expression of the respective ortholog from *S. cerevisiae*. As such, we retrieved the difference, for investigated features, between 1-to-1 orthologous pairs. From analysing *leaky* and *non-leaky* sets within same-species analyses, we find that TR-rate correlates most strongly with gene expression, codon usage (CAI), translational efficiency, whereas only in *S. cerevisiae* does TR correlate with GC-content (see Fig S1 and Table S2). We also find that these features disappear when comparing *leaky* and *non-leaky* orthologs, but that 3’-UTR GC-content remains as a significantly differentiated feature between sets (see Fig S2, Table S4 and S5). Comparisons of *leaky* and *non-leaky* genes - both within same-species, and between orthologs - support the notion that *leaky* genes have an overall higher GC-content of the 3’-UTR than *non-leaky* genes (see Fig S3). These results begs the question if and how the GC-content of the 3’-UTR is associated to TR.

The connection between expression and GC-content could be indication of mRNA stability, which was previously explored in the context of TR, but without verification [9].

High GC-content in the 3’-UTR would by chance lower the probability of random stop codons, and allow translation to continue once the initial stop codon is initially bypassed. In most species, the mutational bias tend to be GC sites that mutate into to TA sites [16]. However, Long et al. [16] found that there is also a substantial contribution to GC composition by either natural selection or biased gene conversion, possibly both. Biased gene conversion that elevates GC-content has previously been suggested to drive emergence of evolutionary novelty in yeasts [17]. However, according to a study that investigated mutation bias in *S. pombe* and *S. cerevisiae*, biased gene conversion does not contribute to the nucleotide composition in *S. pombe* [18]. In light of this, we can not exclude that the difference in 3’-UTR GC-content - between *leaky* and *non-leaky* proteins - may be a result of selection. However, it is ambiguous whether GC-content of the 3’-UTR would be a cause or consequence of TR. There is an upper and lower threshold for the tolerated load of phenotypic mutations, where the lower threshold is suggested to be regulated by translational cost-efficiency [1]. Prior to eliminating mutations, selection may primarily act to minimize deleterious effects of phenotypic mutations, as previously found by [5]. Codons that are GC-rich tend to yield amino acids that are disordered [19] and less prone to aggregate, which may make the extended isoform by TR effectively neutral [9]. Thus, selecting for GC-rich 3’-UTR, yielding GC-rich codons, may buffer potentially deleterious effects of TR by yielding neutral extensions. However, we are unable to test this as one can not infer dN/dS on non-coding regions directly. Further methodological advancement is needed to infer selection on non-coding regions like 3’-UTRs. Alternatively to TR yielding selection upon the 3’-UTR, one may speculate whether elevated GC-content may cause TR. However, to our knowledge, there is no indication of previous research that suggests that GC-content affects ribosomal accuracy. In conclusion, we can not exclude that the GC-content of the 3’-UTR are under selection in *leaky* genes, which may be a consequence of TR. The GC-content of 3’-UTRs should be considered for future investigation regarding TR.

### Evolutionary rate for *S. cerevisiae*

The look ahead effect hypothesis predicts that phenotypic mutations may result in proteins having an elevated evolutionary rate, given that the phenotypic mutations are beneficial. Elevated evolutionary rates would imply that proteins - that endure beneficial phenotypic mutations - undergo positive selection. If TR facilitates the evolution of beneficial gene variants, in accordance with the look-ahead effect [4], we expect that a larger proportion of *leaky* proteins - than *non-leaky* proteins - should be under positive selection. Alternatively, if TR does not yield intermediate phenotypic stepping stones that enhance protein evolvability, we should see no difference between *leaky* and *non-leaky* proteins. Our null hypothesis postulates that proteins do not differ in evolutionary rate whether they are *leaky* or *non-leaky*. To test this, we conducted two inferences of McDonald-Kreitman test (MKT) [20] that infers whether proteins are undergoing purifying selection (see Materials and Methods). We analysed *leaky* and *non-leaky* proteins from *S. cerevisiae*, and compared the two sets for selective pressure. We inferred polymorphism data for *S. cerevisiae* by making use of publicly available data [21] (1102 yeast project). Our focal gene was the lab strain of *S. cerevisiae*, and *S. paradoxus* was used as an outgroup to infer diverging polymorphism (see Materials and Methods).

Initially, we had 80 genes of the *leaky* set and 3200 genes of the *non-leaky* set. After excluding some proteins due to insufficient annotation (see Materials and Methods), we retained 64 *leaky* genes. We found that the majority of *leaky* proteins undergo purifying selection, but so were genes from the *non-leaky* set (see Fig 1). Moreover, the fraction of proteins undergoing positive selection did not alter between the two sets (see Table 1), and we can not reject the null hypothesis. We investigated if any features, beside TR-rate, correlated with purifying selection. We find that gene expression is the strongest correlating factor of evolutionary rate (see Fig S4), which is in accordance with previous studies [22–24]. When we exclusively investigated the *leaky* set (see Fig S5), we saw the same trend. Complete data for the MKT can be found in S2File.txt.

**Fig 1.**
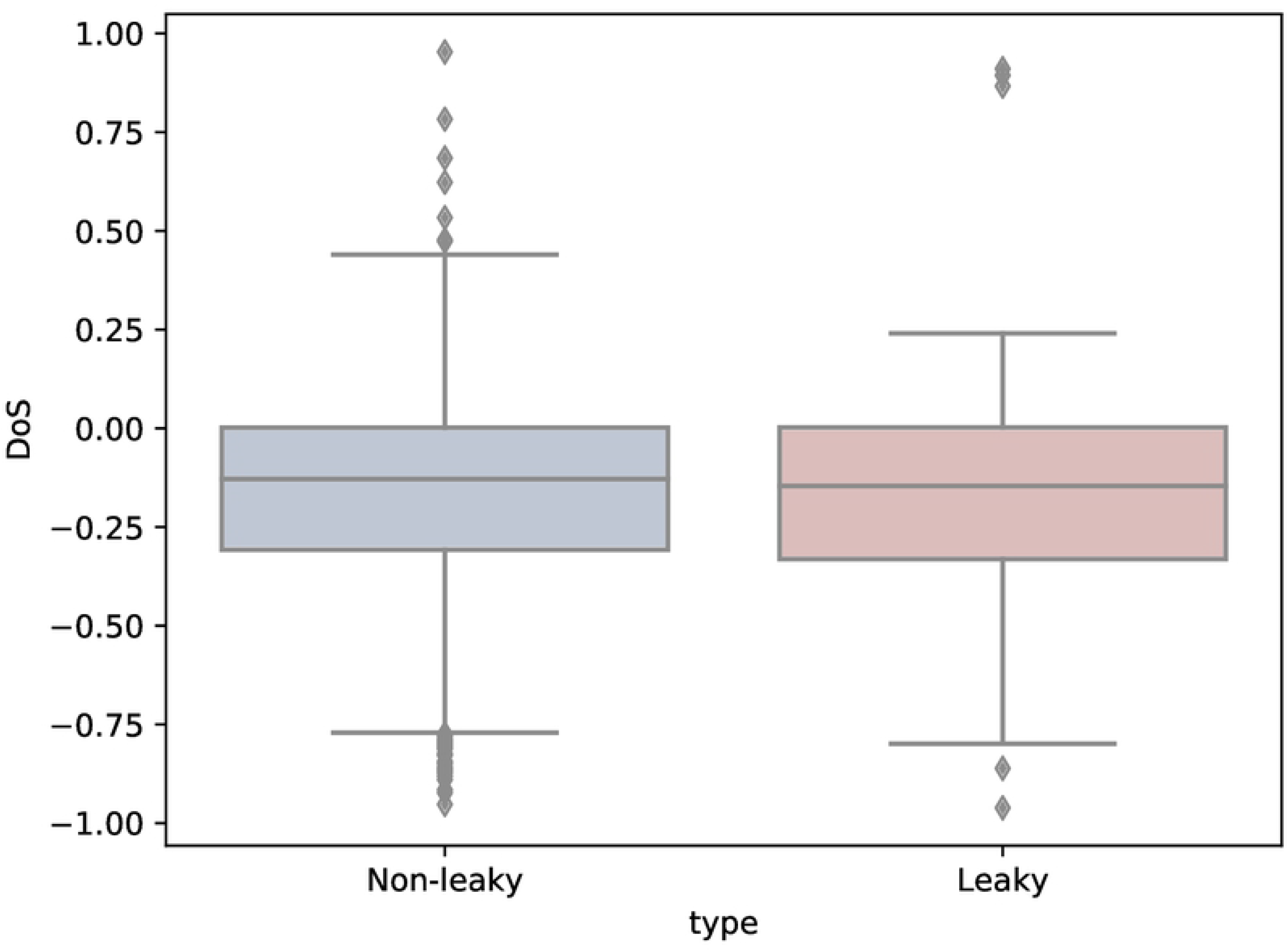
Boxplot of Direction of Selection (DoS) for *leaky* and *non-leaky* sets. Blue depicts the *non-leaky* set and red depicts the *leaky* set. Y-axis depicts direction of selection (DoS). Above 0 indicates that the proteins are under positive selection, and below 0 indicates that the proteins are under purifying selection. Values around 0 indicate neutral evolution.

**Table 1.** Quantity of proteins under different selective pressure, retrieved from McDonald-Kreitman test.

| Set | Positive | Purifying | Neutral |
| --- | --- | --- | --- |
| *leaky* | 7 | 37 | 20 |
| *non-leaky* | 398 | 1868 | 934 |
| *leaky* % | 11% | 58% | 31% |
| *non-leaky* % | 12% | 58% | 29% |

Whether a protein undergoes neutral, positive, or purifying selection, seems to be unrelated to TR. As TR is neither selected against nor for, we find that TR is neutral with respect to selective pressure. However, the fact that TR appears to be neutral may in part be a sampling bias. As we selected *leaky* proteins by genes showing consistent TR - excluding proteins that displayed TR once - we automatically exclude strongly deleterious phenotypic mutations that would already be purged. Thus, including proteins that consistently undergo phenotypic mutations is to include neutral or near-neutral phenotypic mutations. On the other hand, proteins that undergo TR are under both neutral, purifying, and positive selection, which entails that TR prevails in proteins across a range of selective pressures. This suggests that the TR-extensions of *leaky* proteins are sufficiently neutral to not differentiate selection between *leaky* and *non-leaky* proteins on nucleotide level. In conclusion, our analysis refutes the prediction of the look-ahead effect, that proteins undergoing phenotypic mutations would have a rapid evolutionary rate identified by positive selection. We find that TR is neutral and goes unseen by natural selection.

### Paralogs in *S. cerevisiae*

Beyond inferring differential evolutionary rate by comparing the sets, we speculated that one may see a diverging trend by comparing *leaky* and *non-leaky* homologous pairs. Ohno suggested that gene duplication may enable genes to evolve a new function, by maintaining native function by a gene copy [25]. With two gene copies, one gene copy can be exploratory without the cellular system losing the vital gene function [25]. Since function is strongly related to the structure of the encoded protein, it can also be assumed that the initial protein fold would be maintained in either of the two copies (i.e. paralogs). Accordingly, the *leaky* gene would be free to explore the phenotypic landscape while a *non-leaky* paralog would maintain the biological function. In such a scenario, we hypothesized that a high proportion of *leaky* proteins would have *non-leaky* paralogs. Paralogs were retrieved for the *leaky* and *non-leaky* sets for *S. cerevisiae*, using annotations provided by SGD (see Materials and Methods). From the *leaky* set, 35 genes (39%) have an annotated paralog. In the *non-leaky* set, 594 genes (17%) have an annotated paralog. Amongst the *non-leaky* set, the majority of genes (57%) have a *non-leaky* paralog and only a minority (2%) have a *leaky* paralog (see Table 2). As the majority of paralogs are unassigned for the *leaky* set, it is not possible to deduct whether the *leaky* set comprises of primarily *non-leaky* or *leaky* paralogs. The current data suggest that a higher fraction of paralogous pairs are found within the *non-leaky* set. However, the results are biased by the fact that the *non-leaky* set contain more genes. In conclusion, there is no support for that *leaky* genes would be exploratory by the assurance of a *non-leaky* gene-copy.

**Table 2.** Paralogs of the *non-leaky* and *leaky* sets in *S. cerevisiae*.

| Set | Total | NL Paralog | L Paralog | Unassigned |
| --- | --- | --- | --- | --- |
| *NL* | 594 | 340 | 13 | 241 |
| *L* | 35 | 13 | 0 | 22 |
| *NL* | 100% | 57% | 2% | 41% |
| *L* | 100% | 37% | 0% | 62% |

NL stands for *non-leaky* set, whereas L stands for *leaky* set. Unassigned is where we did not know whether a gene was *leaky* or *non-leaky* as it was not part of our expressed data set.

## Conclusion

In order to infer the evolutionary role of TR, we have here conducted comparative analyses between proteins that undergo TR and canonically translated proteins. Our inter-species comparison revealed that TR is not conserved between orthologs, nor that genes are purged on account of TR. Inference of paralogs in *S. cerevisiae* suggests that TR occurs without assurance of functional maintenance by a *non-leaky* paralog. Moreover, inference of selective pressure in *S. cerevisiae* suggest that there is no difference between *leaky* and *non-leaky* proteins. Our inference of the look-ahead effect, that *leaky* proteins would have an elevated evolutionary rate, was not confirmed. Ultimately, our results suggest that TR is a neutral event that prevails without natural selection acting upon it. Taken together with inference of presence and rate of TR, in *S. cerevisiae* and *S. pombe*, our analyses suggest that TR is continuously present at a low rate and not purged. This is in accordance with other studies that find continuous TR to be heterogeneously present [26].

Recently, TR has been suggested to be non-adaptive as its presence appear stochastic [27]. However, biological heterogeneity and stochastic processes is raw material for novel adaptations [28]. Moreover, the lack of strong adaptation is not an argument against evolutionary relevance. Neutrally evolving proteins have previously been suggested to provide a basis for evolutionary novelty [29]. We suggest that phenotypic mutations like TR are permissive. Genotypic and phenotypic mutations occur at a rate governed by the efficiency of natural selection, formulated as the drift-barrier hypothesis [30]. Any phenotypic mutation is initially caused by imperfect purging by natural selection. However, damage control is necessary once TR is present in the cellular environment. Yet, damage control is not needed if the yielded isoform is neutral or near-neutral, and TR can thus persist. However, deleterious isoforms are less likely to persist, as misfolded proteins are degraded by the proteasome [31]. With respect to the abundant findings of TR, by us and others [7,8,32–34], it is doubtful that TR is deleterious as the proteasome would constantly degrade TR-isoforms, which is doubtful given a high cellular energy cost. Rather than direct purging of deleterious mutations, mitigation of deleterious mutations has been found to be the first evolutionary response [5]. Buffering deleterious isoforms by making them neutral would allow TR to persist and explore functional interactions [9]. We suggest that TR-yielded isoforms persist, either because they are initially neutral or have become neutral by selection to mitigate detrimental effects. Like wallflowers, TR-yielded isoforms go unseen and persist. Moreover, in alternating cellular environments, unseen ‘wallflowers” may become relevant. We found that a subset of TR-isoforms in *S. pombe* are homologous to functional encoded proteins in *S. cerevisiae*. This is in accordance with studies who suggest that TR biologically functional and not harmful [7, 8, 32–35]. Overall transcriptional heterogeneity has been found to be beneficial in stressful condition [36, 37], and suggested to be an adaptive trait under selection [38]. Also protein synthesis has been found to be heterogeneous and noisy [8, 26], but also harmful [39]. However, TR have been found to yield positive fitness effects for microorganisms under stress, and plainly neutral in absence of stress [6, 26]. We suggest that the TR-yielded isoforms enrich the proteomic diversity by offering slight variants of encoded proteins that may are readily available in sudden environmental shifts, which has been suggested previously by others [8, 34]. However, the evolutionary trajectory remains unclear for how phenotypic mutations, like TR, would transcend from noise to become adaptive.

To investigate whether there is an adaptive advantage of TR *overall* and whether noise by phenotypic mutations elevate protein evolvability, we believe one needs to infer the effect over shorter time scale than we have done here. By comparing error prone strains to canonical strains - in both ‘‘stressful” and ‘‘controlled” environments - one should be able to test if phenotypic mutations allows for rapid adaptation.

In conclusion, we find that phenotypic mutations, yielded by TR, do not yield rapid adaptation or increase protein evolvability. We suggest that TR goes unseen by natural selection like permissive “wallflowers”. Given that a subset of TR-isoforms are highly homologous to encoded orthologs, we speculate that TR-isoforms may be and become adaptive. However, expanded methodology is needed to infer whether 3’-UTR undergo selection for error mitigation as a consequence of TR. Future studies should also infer whether phenotypic mutations are adaptive by experimental inference, but on short evolutionary time frames.

## Materials and Methods

### Handling of ribosome profiling data

The ribosome profiling data for *S. cerevisiae* were obtained from [10]. Data was obtained and is accessible at NCBI GEO database with accession ID GSE52119, where individual accession files have IDs; SRR948553,SRR948555,SRR948552 and SRR948551. Ribosome profiling data for *S. pombe* was obtained from [11], and was accessed at NCBI GEO database by accession ID GSE98934. Individual files have accession IDs; SRR5564114 and SRR5564124.

Prior to alignment, the reads were trimmed and aligned to rRNA. Reads that did not align to rRNA, were aligned to the genome with bowtie [40] (-S -y -a -m 1 –best –strata -p 22). When aligning reads to the 3’-UTR, we used extra stringent alignment where multimapping was not allowed and only one mismatch was allowed (-S -y -a -m 1 –best –strata -n 0 -e 1 -p 22). To sort the output we used samtools [41, 42].

Genome for *Sacharomyces cerevisiae* S288C was downloaded from Saccharomyces Genome Database (SGD) with annotations [43, 44] on February 8th 2018.

### Detecting translational readthrough

Only expressed genes that have annotated 3’-UTR were included in further analyses. Annotations by Yassour et al. [45] were used for mapping reads to the 3’-UTR. HTSeq was used [46] to retrieve the count number of reads mapped with genes and respective 3’-UTR, using strict-mode that excludes overlapping reads. Genes that consistently were showing translational readthrough (TR) in all replicates were grouped as “*leaky* genes”. Genes displaying TR in some but not all replicates were grouped as “*semi-leaky* genes”. Genes with annotated 3’-UTR without any count hits, consistently between replicates, were grouped as “*non-leaky* genes”.

Mapped reads to 3’-UTR can indicate continued translation of the mRNA beyond the first stop codon, but these reads can also be mere noise. Several measures were made to ensure reads mapped to the 3’-UTR were justifiably counted as translational readthrough (TR). Firstly, annotated 3’-UTRs that are overlapping with a gene on the same strand were excluded. 3’-UTRs with a sequence length shorter than 30 nucleotides were excluded as they infer high stochasticity when calculating coverage.

Before TR rate was estimated, an initial threshold was set for at least 5 reads to be registered as mapped to the 3’-UTR for each replicate. This is a common lower threshold when considering gene expression [47]. TR rate was calculated as the following: The sequence hit count (obtained by HTSeq) was normalised by dividing read length with the sequence length, as done by [48]. The normalized hit count for the 3’-UTR was divided with the normalized hit count value of the protein coding sequence (CDS), yielding relative expression of 3’-UTR. Genes displaying spurious translation by relative expression of one or above were excluded. Relative expression over or near one, effectively implies that the 3’-UTR is being expressed as high as the CDS.

Assuring that the reads were accurately indicating TR, we controlled for background noise and that the TR followed the appropriate open reading frame (ORF). We estimated background noise by quantifying the coverage of riboreads that aligned to tRNA, that were aligned by the same stringent criteria as 3’-UTR. tRNA is not translated by the ribosome. We therefore interpret riboreads aligned to tRNA as noise - either caused by ribosomes that spuriously land on RNA or imperfect alignment. By dividing the read count with sequence length we retrieved the normalised coverage for tRNAs. The highest value - between the replicates, not the mean of the replicates - was used as a threshold for noise: all genes that had a read coverage in the 3’-UTR equal or lower to the tRNA coverage (our threshold) were excluded from our analyses. Lastly, we control for that our indicated TR follow the appropriate open reading frame (ORF). We controlled, by an in house script, that the reads aligned with the ORF up until next stop codon in frame in the 3’-UTR. If the coverage was higher or equal beyond the first stop codon encountered in the 3’-UTR, they were dismissed from further analyses as ambiguous.

### Protein feature analyses

For each replicate of both footprints and RNA-seq, gene expression was calculated as Transcript Per Million (TPM) and then checked for significant distribution differences by a Kolmogorov-Smirnov test, which was non-significant. Translational efficiency (TE) was calculated as described by Ingolia et al. [49], dividing TPM of the ribosome profiling reads by the TPM of the RNA-seq reads. Sequence length was measured in nucleotides of the CDS (not including UTR).

We used the IUPred short algorithm to predict intrinsic disorder in the protein sequences based on the frequency of disorder-promoting amino acids [50], which uses 0.5 as the threshold for a sequence to be disordered.

We analysed codon usage for the proteins within all three sets. We made use of Codon Adaptation index (CAI) [51]. We used codonW version 1.4.2 (http://codonw.sourceforge.net/) to conduct the analyses. We analysed all CDS in all three sets. Moreover, we analysed the last 30 nucleotides of each CDS in all sets as an estimation of the C-termini.

### McDonald-Kreitman test analysis

We inferred polymorphism data for *S. cerevisiae* by making use of publicly available data from [21] (1102 yeast project). The allele sequences were aligned using mafft v7.397 [52, 53]. By an in-house script using Python 2.3, we retrieved synonymous (Ps) or non-synonymous (Pn) codon substitutions within-species. Aligning focal gene of *S. cerevisiae* with ortholog from *S. paradoxus*, we then inferred synonymous (Ds) and non-synonymous (Dn) substitutions. Data for *S. paradoxus* were retrieved from [54]. Our focal gene was the lab strain YGD of *S. cerevisiae*, with annotations from Saccharomyces Genome Database (SGD) [44].

Some of the sequences from the 1102 yeast project have ambiguous nucelotides (eg X instead of A,G,T,C) and inferring synonymous from non-synonymous substitutions was therefore not possible. These sequences were therefore excluded from further analyses, which diminished our data set.

We applied two McDonald-Kreitman tests. An extension of the original McDonald-Kreitman test [20] is inference of the proportion of residues under selection, known as alpha [55]. We applied Fisher’s exact test to infer significance of alpha. The significant data is what we report on with respect to protein undergoing ‘neutral’,’negative’, or ‘positive’ selection. Another inference of McDonald-Kreitman test, which is more robust to sampling bias, is Direction of Selection (DoS) test [56]. Both of these tests indicate that the sets are not different with respect to selection pressure. The resulting values from these analyses can be found in S1File.csv.

### Alignments of orthologs

We used curated annotation of orthologues between *S. cerevisiae* and *S. pombe* [57], retrieved from PomBase [58]. We aligned the *leaky* proteins in *S. pombe* with their*non-leaky* orthologue in *S. cerevisiae*, by using exonerate version 2.4.7 [12]. We used the protein2dna model, using “exhaustive” and “bestfit” parameters. All other parameters were set to default. The *non-leaky* protein sequence from *S. cerevisiae* was used as query sequence, and respective orthologue nucleotide sequence from *S. pombe* was target sequence.

### Retrieval of paralogs

We accessed curated paralogs by the Saccharomyces Genome Database [43] by using yeastmine [59]. Accessing yeastmine by intermine [60, 61], the paralogs of proteins in the *leaky* and *non-leaky* sets were retrieved.

## Supporting information

### Comparison between orthologs

**Table S1.** Conservation of TR between orthologs in *S. pombe* and *S. cerevisiae*.

| *S.pombe* \ *S.cerevisiae* | 1 NL | >1 NL | ≥ 1 NL | ≥ 1 L |
| --- | --- | --- | --- | --- |
| *leaky* | 35% | 23% | 58% | 3% |
| *non – leaky* | 42% | 22% | 64% | 1% |

Percentage of *leaky* and *non-leaky* proteins in *S.pombe* that have orthologs in *S.cerevisiae* that are either *leaky* (L) or *non-leaky* (NL). For example, 35% of *leaky* proteins of *S.pombe* have at least one *non-leaky* ortholog, while 23% of *leaky S.pombe* proteins have more than one *non-leaky* ortholog in *S.cerevisiae*. In total, 58% of *leaky* proteins in *S.pombe* one or more *non-leaky* orthologs, while only three percent have a *leaky* ortholog in *S.cerevisiae*.

### Protein feature analysis

**Table S2.** Spearman rank correlation between variables and TR-rate for *S. cerevisiae* and *S. pombe*.

| Data for *S.pombe* |  |  |  |  | Data for *S.cerevisiae* |  |  |  |
| --- | --- | --- | --- | --- | --- | --- | --- | --- |
| Variables | rho | p | rho* | p* | rho | p | rho* | p* |
| *TPM* | 0.09 | 0.0 | -0.818 | 0.0 | 0.16 | 0.0 | -0.949 | 0.0 |
| *GC 3' – UTR* | 0.21 | 0.0 | 0.03 | 0.64 | 0.03 | 0.09 | 0.69 | 0.0 |
| *GC CDS* | 0.07 | 0.0002 | -0.53 | 0.0 | 0.09 | 0.0 | -0.36 | 0.001 |
| *length* | -0.06 | 0.001 | -0.10 | 0.14 | -0.11 | 0.0 | 0.19 | 0.07 |
| *disorder* | 0.02 | 0.36 | 0.013 | 0.85 | 0.03 | 0.06 | -0.12 | 0.27 |
| *disorder C – termini* | 0.01 | 0.66 | 0.002 | 0.97 | 0.02 | 0.37 | -0.14 | 0.20 |
| *CAI* | 0.038 | 0.04 | -0.55 | 0.0 | 0.13 | 0.0 | -0.89 | 0.0 |
| *CAI end* | 0.016 | 0.38 | -0.21 | 0.002 | 0.11 | 0.0 | -0.73 | 0.0 |
| *TE* | 0.088 | 0.0 | -0.55 | 0.0 | 0.14 | 0.0 | -0.59 | 0.0 |

**Fig S1. Analyses between *leaky* and *non-leaky* sets for respective species.** Unless stated otherwise, the features are data for the CDS or full protein sequence. ‘TR rate’ stands for translational readthrough rate, ‘TPM’ (Transcripts per Million) stands for gene expression, ‘GC’ stands for GC-content, ‘disorder’ stands for intrinsic disorder, ‘TE’ stands for Translational Efficiency, ‘CAI’ stands for Codon Adaptation Index and ‘CAI end’ represents CAI data only for the last 30 nucleotides of the CDS.

**Table S3.**
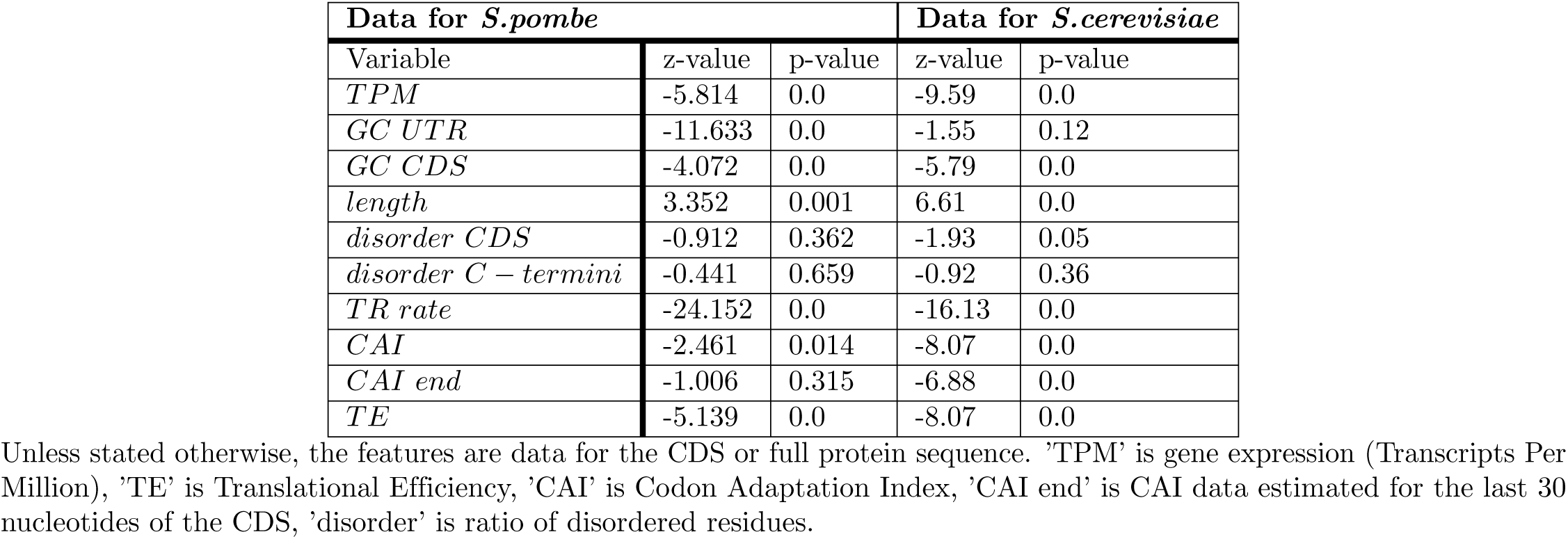
Wilcoxon rank sum test between *leaky* and *non-leaky* sets for features,for *S. cerevisiae* and *S. pombe*.

**Table S4.**
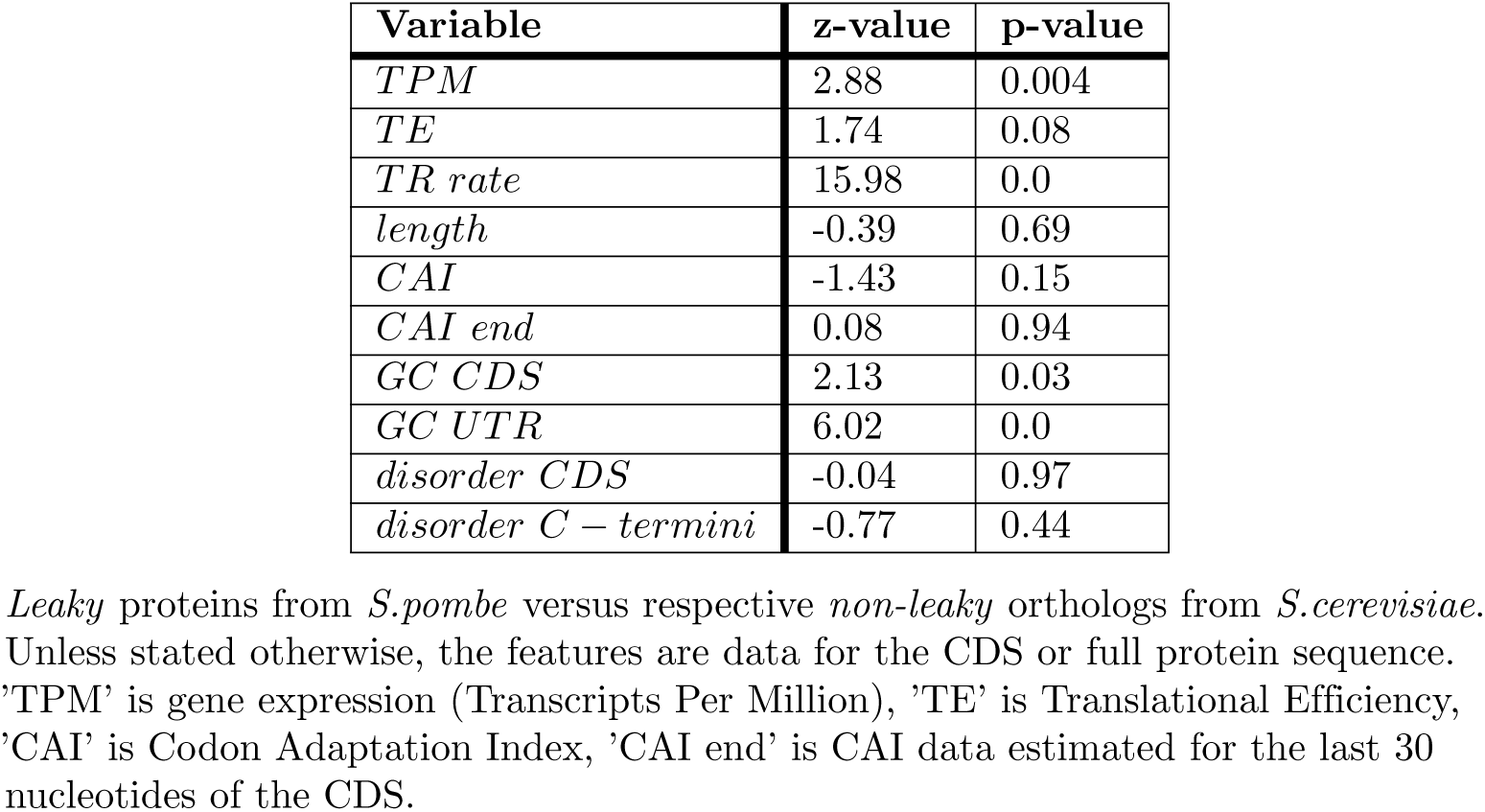
Mann-Whitney Wilcoxon test for variables of orthologous pairs.

| Variable | z-value | p-value |
| --- | --- | --- |
| *TPM* | 2.88 | 0.004 |
| *TE* | 1.74 | 0.08 |
| *TR rate* | 15.98 | 0.0 |
| *length* | -0.39 | 0.69 |
| *CAI* | -1.43 | 0.15 |
| *CAI end* | 0.08 | 0.94 |
| *GC CDS* | 2.13 | 0.03 |
| *GC UTR* | 6.02 | 0.0 |
| *disorder CDS* | -0.04 | 0.97 |
| *disorder C – termini* | -0.77 | 0.44 |

*Leaky* proteins from *S.pombe* versus respective *non-leaky* orthologs from *S.cerevisiae*. Unless stated otherwise, the features are data for the CDS or full protein sequence. 'TPM' is gene expression (Transcripts Per Million), 'TE' is Translational Efficiency, 'CAI' is Codon Adaptation Index, 'CAI end' is CAI data estimated for the last 30 nucleotides of the CDS.

**Fig S2. Difference of features between orthologs, sets shown in different colour.** Difference between *leaky* proteins in *S. pombe* and their *non-leaky* orthologs from *S. cerevisiae* are shown in red. Green display difference between *non-leaky* orthologs between *S. cerevisiae* and *S. pombe*. Unless stated otherwise, the features are data for the CDS or full protein sequence. ‘TPM’ is gene expression (Transcripts Per Million), ‘TE’ is Translational Efficiency, ‘CAI’ is Codon Adaptation Index, ‘CAI end’ is CAI data estimated for the last 30 nucleotides of the CDS.

**Fig S3. GC-content of CDS and 3’-UTR in respective species.** (a) and (b): Distribution of GC-content for leaky and non-leaky sets, where *non-leaky* data is depicted in green and *leaky* data is depicted in red. (c)-(f): Gene expression plotted against GC-content. (g) and (h): TR-rate versus GC-content.

**Table S5.** Spearman rank correlation test of difference of TR-rate and variables.

| Variables | rho | p | rho* | p* |
| --- | --- | --- | --- | --- |
| *TPM* | -0.59 | 0.0 | 0.07 | 0.01 |
| *TE* | -0.29 | 0.01 | 0.04 | 0.10 |
| *length* | -0.36 | 0.001 | -0.01 | 0.61 |
| *CAI* | 0.15 | 0.16 | -0.04 | 0.17 |
| *CAI end* | 0.09 | 0.38 | 0.00 | 0.93 |
| *GC CDS* | -0.19 | 0.08 | 0.06 | 0.04 |
| *GC 3' – UTR* | -0.12 | 0.25 | 0.16 | 0.0 |
| *disorder CDS* | -0.02 | 0.82 | -0.00 | 0.96 |
| *disorder C – term* | 0.01 | 0.94 | -0.02 | 0.44 |

Data include differences between *non-leaky* orthologs and *leaky* versus *non-leaky* orthologs. Asterisk (\*) includes data only for the difference between *leaky* proteins in *S.pombe* and their *S.cerevisiae non-leaky* orthologs. Unless stated otherwise, the features are data for the CDS or full protein sequence. 'TPM' is gene expression (Transcripts Per Million), 'TE' is Translational Efficiency, 'CAI' is Codon Adaptation Index, 'CAI end' is CAI data estimated for the last 30 nucleotides of the CDS.

### Evolutionary analysis

**Fig S4. Selective pressure and protein features for *leaky* set in *S. cerevisiae***. Selection pressure marked by colour for *leaky* proteins in *S. cerevisiae*, retrieved by McDonald-Kreitman test. All values are log transformed. The colour depicts what selective pressure the proteins are under, as depicted by the legend: orange is positive selection, blue is negative (or purifying) selection, whereas green represents unknown selective pressure, or neutral evolution. Axes: ‘TR rate’ is rate of translational readthrough, ‘TPM’ is Transcripts Per Million and infers gene expression. ‘Pn’ and ‘Ps’ is non-synonymous and synonymous polymorphisms within *S. cerevisiae* population data, whereas ‘Ds’ and ‘Dn’ is synonymous and non-synonymous substitutions, inferred between *S. cerevisiae* and *S. paradoxus*.

**Fig S5. Selective pressure and protein features for *leaky* and *non-leaky* sets in *S. cerevisiae*.** Selection pressure marked by colour for both *leaky* and *non-leaky* proteins in *S. cerevisiae*, retrieved by McDonald-Kreitman test. All values are log transformed. The colour depicts what selective pressure the proteins are under, as depicted by the legend: orange is positive selection, blue is negative (or purifying) selection, whereas green represents unknown selective pressure, or neutral evolution. Axes: ‘TR rate’ is rate of translational readthrough, ‘TPM’ is Transcripts Per Million and infers gene expression. ‘Pn’ and ‘Ps’ is non-synonymous and synonymous polymorphisms within *S. cerevisiae* population data, whereas ‘Ds’ and ‘Dn’ is synonymous and non-synonymous substitutions, inferred between *S. cerevisiae* and *S. paradoxus*.

### Supporting Data

**S1 File.txt Ortholog alignments.**

**S2File.txt Data for and from McDonald-Kreitman test**

**S3File.csv Data for sequence feature analysis for *S. pombe***

**S4File.csv Data for sequence feature analysis for S.cerevisiae**.

## Author contributions statement

E.B.B. and A.S.K. conceived the experiments, A.S.K conducted the experiments and analyses. Both authors reviewed the manuscript.

## Additional information

The authors declare no competing interests.

## References

1. Burger R, Willensdorfer M, Nowak MA. Why are phenotypic mutation rates much higher than genotypic mutation rates? Genetics. 2006;172(1):197–206.

2. Lynch M. The cellular, developmental and population-genetic determinants of mutation-rate evolution. Genetics. 2008;180(2):933–943.

3. Kramer EB, Farabaugh PJ. The frequency of translational misreading errors in E. coli is largely determined by tRNA competition. RNA. 2007;13(1):87–96.

4. Whitehead DJ, Wilke CO, Vernazobres D, Bornberg-Bauer E. The look-ahead effect of phenotypic mutations. Biol Direct. 2008;3:18.

5. Bratulic S, Gerber F, Wagner A. Mistranslation drives the evolution of robustness in TEM-1 Î^2^-lactamase. Proc Natl Acad Sci USA. 2015;112(41):12758–12763.

6. Yanagida H, Gispan A, Kadouri N, Rozen S, Sharon M, Barkai N, et al. The Evolutionary Potential of Phenotypic Mutations. PLoS Genet. 2015;11(8):e1005445.

7. Freitag J, Ast J, Bolker M. Cryptic peroxisomal targeting via alternative splicing and stop codon read-through in fungi. Nature. 2012;485(7399):522–525.

8. Dunn JG, Foo CK, Belletier NG, Gavis ER, Weissman JS. Ribosome profiling reveals pervasive and regulated stop codon readthrough in Drosophila melanogaster. Elife. 2013;2:e01179.

9. Kleppe AS, Bornberg-Bauer E. Robustness by intrinsically disordered C-termini and translational readthrough. Nucleic Acids Res. 2018;46(19):10184–10194.

10. Spealman P, Naik AW, May GE, Kuersten S, Freeberg L, Murphy RF, et al. Conserved non-AUG uORFs revealed by a novel regression analysis of ribosome profiling data. Genome Res. 2018;28(2):214–222.

11. Guydosh NR, Kimmig P, Walter P, Green R. Regulated Ire1-dependent mRNA decay requires no-go mRNA degradation to maintain endoplasmic reticulum homeostasis in S. pombe. Elife. 2017;6.

12. Slater GS, Birney E. Automated generation of heuristics for biological sequence comparison. BMC Bioinformatics. 2005;6:31.

13. Zakon HH. Convergent evolution on the molecular level. Brain Behav Evol. 2002;59(5-6):250–261.

14. Parker J, Tsagkogeorga G, Cotton JA, Liu Y, Provero P, Stupka E, et al. Genome-wide signatures of convergent evolution in echolocating mammals. Nature. 2013;502(7470):228–231.

15. Zou Z, Zhang J. No genome-wide protein sequence convergence for echolocation. Mol Biol Evol. 2015;32(5):1237–1241.

16. Long H, Sung W, Kucukyildirim S, Williams E, Miller SF, Guo W, et al. Evolutionary determinants of genome-wide nucleotide composition. Nat Ecol Evol. 2018;2(2):237–240.

17. Vakirlis N, Hebert AS, Opulente DA, Achaz G, Hittinger CT, Fischer G, et al. A Molecular Portrait of De Novo Genes in Yeasts. Mol Biol Evol. 2018;35(3):631–645.

18. Farlow A, Long H, Arnoux S, Sung W, Doak TG, Nordborg M, et al. The Spontaneous Mutation Rate in the Fission Yeast Schizosaccharomyces pombe. Genetics. 2015;201(2):737–744.

19. Basile W, Sachenkova O, Light S, Elofsson A. High GC content causes orphan proteins to be intrinsically disordered. PLoS Comput Biol. 2017;13(3):e1005375.

20. McDonald JH, Kreitman M. Adaptive protein evolution at the Adh locus in Drosophila. Nature. 1991;351(6328):652–654.

21. Peter J, De Chiara M, Friedrich A, Yue JX, Pflieger D, Bergstrom A, et al. Genome evolution across 1,011 Saccharomyces cerevisiae isolates. Nature. 2018;556(7701):339–344.

22. Drummond DA, Bloom JD, Adami C, Wilke CO, Arnold FH. Why highly expressed proteins evolve slowly. Proc Natl Acad Sci USA. 2005;102(40):14338–14343.

23. Drummond DA, Wilke CO. Mistranslation-induced protein misfolding as a dominant constraint on coding-sequence evolution. Cell. 2008;134(2):341–352.

24. Yang Z, Nielsen R. Mutation-selection models of codon substitution and their use to estimate selective strengths on codon usage. Mol Biol Evol. 2008;25(3):568–579.

25. Ohno S. Gene duplication and the uniqueness of vertebrate genomes circa 1970-1999. Semin Cell Dev Biol. 1999;10(5):517–522.

26. Fan Y, Evans CR, Barber KW, Banerjee K, Weiss KJ, Margolin W, et al. Heterogeneity of Stop Codon Readthrough in Single Bacterial Cells and Implications for Population Fitness. Mol Cell. 2017;67(5):826–836.

27. Li C, Zhang J. Stop-codon read-through arises largely from molecular errors and is generally nonadaptive. PLoS Genet. 2019;15(5):e1008141.

28. Tawfik DS. Messy biology and the origins of evolutionary innovations. Nat Chem Biol. 2010;6(10):692–696.

29. Ruiz-Orera J, Verdaguer-Grau P, Villanueva-Canas JL, Messeguer X, Alba MM. Translation of neutrally evolving peptides provides a basis for de novo gene evolution. Nat Ecol Evol. 2018;2(5):890–896.

30. Lynch M. The Origins of Genome Architecture 2007. Science (New York, NY). 2007;302(5649):1401–1404. doi:10.1126/science.1089370.

31. Rousseau A, Bertolotti A. Regulation of proteasome assembly and activity in health and disease. Nat Rev Mol Cell Biol. 2018;19(11):697–712.

32. Loughran G, Chou MY, Ivanov IP, Jungreis I, Kellis M, Kiran AM, et al. Evidence of efficient stop codon readthrough in four mammalian genes. Nucleic Acids Res. 2014;42(14):8928–8938.

33. Stiebler AC, Freitag J, Schink KO, Stehlik T, Tillmann BA, Ast J, et al. Ribosomal readthrough at a short UGA stop codon context triggers dual localization of metabolic enzymes in Fungi and animals. PLoS Genet. 2014;10(10):e1004685.

34. Schueren F, Thoms S. Functional Translational Readthrough: A Systems Biology Perspective. PLoS Genet. 2016;12(8):e1006196.

35. Pancsa R, Macossay-Castillo M, Kosol S, Tompa P. Computational analysis of translational readthrough proteins in Drosophila and yeast reveals parallels to alternative splicing. Sci Rep. 2016;6:32142.

36. Holland SL, Reader T, Dyer PS, Avery SV. Phenotypic heterogeneity is a selected trait in natural yeast populations subject to environmental stress. Environ Microbiol. 2014;16(6):1729–1740.

37. Bodi Z, Farkas Z, Nevozhay D, Kalapis D, Lazar V, Csorg? B, et al. Correction: Phenotypic heterogeneity promotes adaptive evolution. PLoS Biol. 2017;15(6):e1002607.

38. Wolf MY, Wolf YI, Koonin EV. Comparable contributions of structural-functional constraints and expression level to the rate of protein sequence evolution. Biol Direct. 2008;3:40.

39. Evans CR, Fan Y, Weiss K, Ling J. Errors during Gene Expression: Single-Cell Heterogeneity, Stress Resistance, and Microbe-Host Interactions. mBio. 2018;9(4). doi:10.1128/mBio.01018-18.

40. Langmead B, Trapnell C, Pop M, Salzberg SL. Ultrafast and memory-efficient alignment of short DNA sequences to the human genome. Genome Biol. 2009;10(3):R25.

41. Li H, Handsaker B, Wysoker A, Fennell T, Ruan J, Homer N, et al. The Sequence Alignment/Map format and SAMtools. Bioinformatics. 2009;25(16):2078–2079.

42. Li H. A statistical framework for SNP calling, mutation discovery, association mapping and population genetical parameter estimation from sequencing data. Bioinformatics. 2011;27(21):2987–2993.

43. Cherry JM, Hong EL, Amundsen C, Balakrishnan R, Binkley G, Chan ET, et al. Saccharomyces Genome Database: the genomics resource of budding yeast. Nucleic Acids Res. 2012;40(Database issue):D700–705.

44. Engel SR, Dietrich FS, Fisk DG, Binkley G, Balakrishnan R, Costanzo MC, et al. The reference genome sequence of Saccharomyces cerevisiae: then and now. G3 (Bethesda). 2014;4(3):389–398.

45. Yassour M, Pfiffner J, Levin JZ, Adiconis X, Gnirke A, Nusbaum C, et al. Strand-specific RNA sequencing reveals extensive regulated long antisense transcripts that are conserved across yeast species. Genome Biol. 2010;11(8):R87.

46. Anders S, Pyl PT, Huber W. HTSeq–a Python framework to work with high-throughput sequencing data. Bioinformatics. 2015;31(2):166–169.

47. Anders S, McCarthy DJ, Chen Y, Okoniewski M, Smyth GK, Huber W, et al. Count-based differential expression analysis of RNA sequencing data using R and Bioconductor. Nat Protoc. 2013;8(9):1765–1786.

48. Namy O, Hatin I, Rousset JP. Impact of the six nucleotides downstream of the stop codon on translation termination. EMBO Rep. 2001;2(9):787–793.

49. Ingolia NT, Ghaemmaghami S, Newman JR, Weissman JS. Genome-wide analysis in vivo of translation with nucleotide resolution using ribosome profiling. Science. 2009;324(5924):218–223.

50. Dosztanyi Z, Csizmok V, Tompa P, Simon I. IUPred: web server for the prediction of intrinsically unstructured regions of proteins based on estimated energy content. Bioinformatics. 2005;21(16):3433–3434.

51. Sharp PM, Li WH. The codon Adaptation Index–a measure of directional synonymous codon usage bias, and its potential applications. Nucleic Acids Res. 1987;15(3):1281–1295.

52. Katoh K, Standley DM. MAFFT multiple sequence alignment software version 7: improvements in performance and usability. Mol Biol Evol. 2013;30(4):772–780.

53. Katoh K, Misawa K, Kuma K, Miyata T. MAFFT: a novel method for rapid multiple sequence alignment based on fast Fourier transform. Nucleic Acids Res. 2002;30(14):3059–3066.

54. Scannell DR, Zill OA, Rokas A, Payen C, Dunham MJ, Eisen MB, et al. The Awesome Power of Yeast Evolutionary Genetics: New Genome Sequences and Strain Resources for the Saccharomyces sensu stricto Genus. G3 (Bethesda). 2011;1(1):11–25.

55. Eyre-Walker A. The genomic rate of adaptive evolution. Trends Ecol Evol (Amst). 2006;21(10):569–575.

56. Stoletzki N, Eyre-Walker A. Estimation of the neutrality index. Mol Biol Evol. 2011;28(1):63–70.

57. Wood V, Harris MA, McDowall MD, Rutherford K, Vaughan BW, Staines DM, et al. PomBase: a comprehensive online resource for fission yeast. Nucleic Acids Res. 2012;40(Database issue):D695–699.

58. Lock A, Rutherford K, Harris MA, Hayles J, Oliver SG, Bahler J, et al. PomBase 2018: user-driven reimplementation of the fission yeast database provides rapid and intuitive access to diverse, interconnected information. Nucleic Acids Res. 2019;47(D1):D821–D827.

59. Balakrishnan R, Park J, Karra K, Hitz BC, Binkley G, Hong EL, et al. YeastMine–an integrated data warehouse for Saccharomyces cerevisiae data as a multipurpose tool-kit. Database (Oxford). 2012;2012:bar062.

60. Kalderimis A, Lyne R, Butano D, Contrino S, Lyne M, Heimbach J, et al. InterMine: extensive web services for modern biology. Nucleic Acids Res. 2014;42(Web Server issue):W468–472.

61. Smith RN, Aleksic J, Butano D, Carr A, Contrino S, Hu F, et al. InterMine: a flexible data warehouse system for the integration and analysis of heterogeneous biological data. Bioinformatics. 2012;28(23):3163–3165.

